# Prominent elevation of extracellular matrix molecules in intracerebral hemorrhage

**DOI:** 10.1101/2023.05.26.542446

**Authors:** Hongmin Li, Samira Ghorbani, Ruiyi Zhang, Vincent Ebacher, Erin L. Stephenson, V. Wee Yong, Mengzhou Xue

## Abstract

**Background:** Intracerebral hemorrhage (ICH) is the predominant type of hemorrhagic stroke with high mortality and disability. In other neurological conditions, the deposition of extracellular matrix (ECM) molecules is a prominent obstacle for regenerative processes and an enhancer of neuroinflammation. Whether ECM molecules alter in composition after ICH, and which ECM members may inhibit repair, remain unknown in hemorrhagic stroke.

**Methods:** The collagenase-induced ICH mouse model and an autopsied human ICH specimen were investigated for expression of ECM members by immunofluorescence microscopy. Confocal image z-stacks were analyzed with Imaris 3D to assess the association of immune cells and ECM molecules. Sections from a mouse model of multiple sclerosis were used as disease controls. Tissue culture was employed to examine the roles of ECM members on oligodendrocyte precursor cells (OPCs).

**Results:** Amongst the lectican chondroitin sulphate proteoglycan (CSPG) members, neurocan but not aggrecan, versican-V1 and versican-V2 was prominently expressed in perihematomal tissue and lesion core compared to the contralateral area in murine ICH. Fibrinogen, fibronectin and heparan sulphate proteoglycan (HSPG) were also elevated after murine ICH while thrombospondin was not. Confocal microscopy with Imaris 3D rendering co-localized neurocan, fibrinogen, fibronectin and HSPG molecules to Iba1^+^ microglia/macrophages or GFAP^+^ astrocytes. Marked differentiation from the multiple sclerosis model was observed, the latter with high versican-V1 and negligible neurocan. In culture, purified neurocan inhibited adhesion and process outgrowth of OPCs, which are early steps in myelination in vivo. The prominent expression of neurocan in murine ICH was corroborated in human ICH sections.

**Conclusion:** ICH caused distinct alterations in ECM molecules. Amongst CSPG members, neurocan was selectively upregulated in both murine and human ICH. In tissue culture, neurocan impeded the properties of oligodendrocyte lineage cells. Alterations to the ECM in ICH may adversely affect reparative outcomes after stroke.

**What is already known on this topic** – CSPGs are known to be elevated in multiple sclerosis and intraventricular hemorrhage, where they act as inhibitors of repair by hindering remyelination and axonal regeneration, as well as promoting neuroinflammation. However, there is currently no literature available regarding the role of CSPGs in ICH.

**What this study adds** – Our findings demonstrate the elevation of specific ECM molecules, particularly neurocan, in murine and human ICH. These matrix molecules will likely affect subsequent reparative processes such as remyelination, as suggested by the result that purified neurocan impairs the process outgrowth and maturation of oligodendrocyte precursor cells.

**How this study might affect research, practice or policy** – The targeting of ECM molecules represents a promising strategy to promote remyelination and control neuroinflammation, with the potential to improve prognosis following ICH.

## Introduction

ICH is a devastating disorder characterized by formation of hematoma within the brain parenchyma, with possible blood extension into the ventricles.[1, 2] ICH accounts for 15 to 20% of all strokes, with a disability rate of 70% and a mortality rate of over 50% annually.[3] The poor recovery of ICH generally results from the ongoing toxic neuroinflammation following the mechanical disruption of extravasated blood.[4, 5] Recovery may also be affected by brain ECM components that impact inflammation and repair following other types of central nervous system (CNS) injury.[6, 7]

The ECM is an intricate network of proteins and sugars dispersed throughout the extracellular space of the CNS.[8] The ECM can be segregated into the basement membrane around the cerebral microvessels, interstitial matrix between cellular structures in the parenchyma, and perineuronal nets around certain neuronal soma.[9] The major components of the ECM with known roles in injury and repair include chondroitin sulphate proteoglycans (CSPGs), fibrinogen, fibronectin, heparan sulphate proteoglycans (HSPGs), and thrombospondins (TSPs).

CSPGs are widely expressed in the CNS. They consist of a central core protein and varying number of glycosaminoglycan (GAG) side chains.[8] The lectican CSPG subfamily include aggrecan, brevican, neurocan and versican.[10] Versicans have five splice isoforms: V0 (containing both GAG-beta and alpha domains), V1 (only GAG-beta domain), V2 (only GAG-alpha domain), V3 (neither GAG-beta nor alpha domains) and V4 (part of GAG-beta domain).[11] Fibrinogen, a blood-derived soluble protein synthesized by hepatocytes[12] enters the CNS parenchyma and is converted into insoluble fibrin in CNS injury with blood-brain barrier disruption.[13] Fibronectin exists as a protein dimer through two disulfide bridges and acts as a scaffold for adhering cells, binding heparan and collagen.[14] Heparan sulphate proteoglycans (HSPGs) are found in basement membrane, and are comprised of a core protein with attached heparan sulfate GAGs.[15] Thrombospondins (TSPs) are a family of calcium-binding ECM glycoproteins that regulate cell-cell and cell-matrix interactions.[16]

The ECM is essential for homeostasis and serves metabolic and structural functions in the CNS. However, ECM components are deposited and remodeled aberrantly in neurological diseases and this affects both injury and repair.[8] For example, various ECM members including versican-V1, fibrinogen, fibronectin, HSPGs and TSP are elevated at the peak of clinical disability in an animal model of multiple sclerosis.[7, 17, 18] Sources of ECM deposition after injury include astrocytes, oligodendrocytes, neurons, pericytes and endothelial cells.[17] In chronic multiple sclerosis lesions, the expression of lectican CSPGs including aggrecan and versican rises.[6, 19] Following lysolecithin-induced demyelination, another model of multiple sclerosis, the level of versican-V1 is elevated in lesions.[20]

There is currently a gap of knowledge on the expression and functions of ECM components in ICH. After intraventricular hemorrhage in premature rabbit pups, levels of CSPGs are elevated in the forebrain.[21] As pro-inflammatory macrophages can produce versican-V1,[17, 22] and as chronic CSPG deposition inhibits repair processes after spinal cord injury,[23, 24] it is worth considering whether particular CSPG members are elevated after ICH to impede attempts at regeneration. Moreover, as versican-V1 promotes cytotoxic neuroinflammation in a model of multiple sclerosis, [7] and CSPGs enhance the production of ICH-associated matrix metalloproteinases (MMPs) and pro-inflammatory cytokines,[25] it is imperative to determine whether the ECM is dysregulated in ICH.

In this study, we examined whether and which ECM components are altered after ICH, with focus in the perihematomal area and lesion core. We determined which cell type expresses the elevated ECM molecules, and whether a particular ECM molecule is potentially inhibitory to OPCs in ICH. These results were contrasted with specimens from experimental autoimmune encephalomyelitis (EAE), an animal model of multiple sclerosis used as positive controls for antibody staining. Our overall objective is to derive new insights in whether the ECM is a novel target to improve prognosis after ICH.

## Methods

All animal experiments were performed with ethics approval (protocol number AC21-0073) from the Animal Care Committee at the University of Calgary under regulations of the Canadian Council of Animal Care.

### Mice

Male C57BL/6 wildtype mice (8-12 weeks old) for ICH were purchased from Charles River. C57BL/6 wildtype mice (8–12 weeks old) for EAE were acquired from Jackson Laboratories. Litters from pregnant CD1 mice (P1-P2) were purchased from Charles River and used for OPC cultures. Mice were housed at 22°C, with 12h light and 12h dark cycle, environmental enrichment and free access to food and water.

### ICH induction

The protocol for induction has been described elsewhere.[26] In brief, 0.05U of collagenase type VII dissolved in 0.5μl of saline was injected at a rate of 0.1μl/min over 5 minutes into the right striatum. Then, the needle was maintained at the same spot for additional 5 minutes to prevent reflux. The mice were sutured and then monitored in a thermally controlled environment until recovery.

### EAE induction and tissue harvest

Mice were injected subcutaneously with 200 μg MOG35-55 peptide emulsified in complete Freund’s adjuvant containing 10 mg/ml of heat inactivated Mycobacterium tuberculosis H37RA as described elsewhere.[7] On the day of immunization and 48h later, animals received intraperitoneal injections of 300 ng of pertussis toxin (List Biological Laboratories). Spinal cord sections from mice killed at peak clinical severity (around day 18 after MOG) were used for immunofluorescence analyses.

### ICH brain tissue harvest

Mice were sacrificed at 7-days post-collagenase injection with a lethal dose of ketamine and xylazine. Animals were perfused with a total of 10ml of phosphate-buffered saline (PBS) and 10ml of 4% paraformaldehyde (PFA) in PBS via cardiac puncture. The whole brain was collected into 4% PFA in PBS for fixation overnight, and then was transferred into 30% sucrose solution for 72h. The cerebellum was excised, and the remaining brain tissue was frozen in FSC 22 frozen section media (Leica). Brain blocks were cut coronally by a cryostat into 20 μm sections, collected onto microscope slides and stored at -20°C before staining.

### Human ICH specimen

Paraffin-embedded blocks from an autopsied ICH subject was from Foothills Medical Center, University of Calgary. The subject was a 70-year-old male who had a right middle cerebral artery (MCA) infarct 5 days before death, and with a hemorrhagic transformation 2 days after. Thus, death occurred 3 days after ICH. Sample was collected with full informed consent for autopsy and retention of tissues for research. Gross examination revealed a substantial hemorrhage in the right cerebral parenchyma, along with edema and gyral flattening. The perihematomal regions (because the main hemorrhage region was purely hemorrhage and necrotic tissue) and the contralateral side containing both frontal white matter and cortical regions were taken for research. The use of these human tissues in Calgary for research was approved by the Conjoint Health Research Ethics Board at the University of Calgary (Ethics ID REB15-0444).

### Antibodies

The primary and secondary antibodies used are displayed in Table 1.

**Table 1:** Primary and secondary antibodies.

| Target | Antibody | Commercial source | Catalog number | Dilution from supplier stock |
| --- | --- | --- | --- | --- |
| Astrocytes | Chicken anti-mouse glial fibrillary acidic protein (GFAP) | BioLegend | 829401 | 1:1000 |
| Microglia/macrophages | Goat anti-human/mouse Iba1 | ThermoFisher | PA5-18039 | 1:250 |
| Neuron | Rabbit anti-human/mouse NeuN | Abcam | AB 177487 | 1:500 |
| Versican-V1 | Rabbit anti-mouse versican V0/V1 | Millipore | AB 1033 | 1:100 |
| Versican-V2 | Rabbit anti-mouse versican V0/V2 | Millipore | AB 1032 | 1:100 |
| Neurocan | Rabbit anti-mouse neurocan | Millipore | ABT 1347 | 1:100 |
| Fibrinogen | Rabbit anti-human/mouse fibrinogen | Abcam | AB 34269 | 1:100 |
| Fibronectin | Rabbit anti-human/mouse fibronectin | Abcam | AB 23750 | 1:100 |
| Thrombospondin-1 | Rabbit anti-human/mouse thrombospondin-1 | Abcam | AB 85762 | 1:100 |
| Aggrecan | Rabbit anti-mouse aggrecan | Millipore | AB 1031 | 1:100 |
| HSPG | Mouse anti-mouse heparan sulfate proteoglycan | Amsbio | 370255 | 1:100 |
| CSPG | Mouse anti-mouse CSPG | Millipore | MAB2030 | 1:100 |
| Oligodendrocyte | Mouse anti-human/mouse oligodendrocyte marker O4 | R&D | MAB1326 | 1:50 |
| Isotype | Rabbit isotype control | Abcam | AB 37415 | 1:100 |
| Nuclear | DAPI | ThermoFisher | 62248 | 1:800 |
| Rabbit IgG | Alexa Fluor 647 donkey anti-rabbit IgG | Jackson ImmunoResearch | 711-605-152 | 1:400 |
| Goat IgG | Alexa Fluor 488 donkey anti-goat IgG | Jackson ImmunoResearch | 705-545-147 | 1:400 |
| Mouse IgM | Alexa Fluor 488 donkey anti-mouse IgM | Jackson ImmunoResearch | 715-545-140 | 1:400 |
| Chicken IgY | Cyanine Cy3 donkey anti-chicken IgY | Jackson ImmunoResearch | 703-165-155 | 1:400 |
| Goat IgG | Cyanine Cy3 donkey anti-goat IgG | Jackson ImmunoResearch | 705-165-147 | 1:400 |

### Immunofluorescence staining

Microscope slides containing mouse brain tissues were thawed at room temperature for 30 minutes, then hydrated with PBS for 5 minutes, and permeabilized with 0.1% Triton X-100 in PBS for 5 minutes. Tissue sections were blocked by horse serum blocking solution (0.01 M PBS, 1% bovine serum albumin (BSA), 10% horse serum, 0.1% Triton-X100, 0.1% cold fish gelatin, and 0.05% Tween-20) for 1h. Alternatively, for staining using the HSPG antibody in mice, AffiniPure Fab fragment donkey anti-mouse IgG (H+L) (Jackson ImmunoResearch, 715-007-003, 1:50) was added to the blocking buffer. Tissues were incubated with primary antibodies suspended in antibody dilution buffer (0.01 M PBS, 1% BSA, 0.1% Triton-X100, 0.1% cold fish gelatin) overnight at 4°C. Next, slides were washed with PBS containing 0.2% Tween-20 and incubated with fluorophore conjugated secondary antibodies (1:400) and 1 μg/ml of DAPI for 1h. The slides were washed and mounted using Fluoromount-G solution (SouthernBiotech).

For CSPG staining, chondroitinase ABC (ChABC, Sigma) digestion was performed prior to the blocking step in order to remove GAG chains to facilitate antibody binding to the core protein. Slides were then incubated with ChABC diluted in PBS (0.2 U/mL) for 30 minutes at 37°C.

For human paraffin sections, these were cut (7 μm) using a Leica RM2135 Microtome. Following deparaffinization, sections were subjected to antigen retrieval by boiling in 10 mM sodium citrate buffer (pH 6.0) for 20 minutes, then enzymatically digested with ChABC at 37°C for 30 minutes. The sections were next incubated with 4% horse serum to block nonspecific binding, then incubated with anti neurocan, anti-GFAP and anti-Iba1 overnight at 4°C. Secondary antibodies (Table 1) were added for 60 minutes. Fluorescence images were taken on a confocal microscope (Fluoview FV10i; Olympus).

### Cell culture

#### Mouse oligodendrocyte precursor cells (OPC)

Brains from postnatal day P0-2 mouse pups were isolated and processed for OPCs as described previously.[7] OPCs were seeded at a density of 1 × 10^4^ cells per well in 100 μL of oligodendrocyte differentiating medium (described below) and grown at 37°C and 8.5% CO2 for 24–72 h. The coating of these 96-well flat bottom black plates was either poly-L-lysine (10 μg/ml, control), neurocan (10 μg/ml, Millipore), a mixed CSPG preparation (10 μg/ml, Millipore), fibronectin (10 μg/ml, Sigma) or fibrinogen (10 μg/ml, Millipore). Culture medium is described elsewhere.[20] The purity of olig2+ oligodendrocyte lineage cells was over 80%.

For immunocytochemistry, OPCs were fixed for 10 min at RT using 4% paraformaldehyde, rinsed with PBS and then permeabilized with 0.2% Triton X-100 for 10 min at room temperature. The primary antibody mouse anti-mouse sulfatide O4 (oligodendrocyte lineage cell marker) (R&D) was applied overnight at 4°C, followed by secondary antibody and DAPI. Cells were analysed with a Molecular Devices ImageXpress Micro XLS.[20]

#### Widefield and confocal fluorescence microscopy

Overviews of brain sections were imaged with an Olympus VS120 slide scanner using a 10×/0.4NA objective. These were used to locate the lesions. Only the brightness and contrast were adjusted to better display representative images. All samples were then imaged on a Leica TCS SP8 laser scanning confocal microscope using a 25×/0.95 NA water objective. Three-dimensional (3D) z-stacks images (2048 × 2048 × 37 voxels) of the four fluorescent probes were acquired using the 405 nm, 488 nm, 552 nm, and 640 nm lasers sequentially at either 228 × 228 × 567 nm or 114 × 114 × 567 nm voxel size. The two different resolutions were used for quantifying % area of ECM molecules in Regions of Interest (ROIs) and % positive cells with ECM molecules, respectively. Imaging parameters were kept constant for each set of experiments.

#### Confocal image analysis

ImageJ software (NIH) was used to quantify the % ECM molecules in ROIs. For each z-stack image, maximum-intensity projections were created and ROIs were drawn around the perihematomal, lesion center, and contralateral areas according to GFAP or Iba1 labeling. Image segmentation was achieved using a set intensity threshold for each probe. Negative secondary antibody controls or contralateral controls were used to assess baseline signals and determine thresholds. For each probe, intensity threshold as well as size and circularity filters were kept constant across all samples for each experimental set. Total area and percent area (i.e., % ECM molecules in ROI) were measured.

Imaris software (Oxford Instruments) was used to determine the percentage of Iba1^+^ or GFAP^+^ cells overlapping with ECM molecules. Labelled areas were segmented as surfaces via a set intensity threshold determined as above. Iba1^+^ or GFAP^+^ cells were separated using seed points within the Imaris Surface creation workflow to get the total number of cells. For each probe, intensity threshold, surface details were kept constant within each set of experiments. % positive cells with ECM molecules were obtained by dividing the number of positive cells with ECM molecule by the total number of positive cells.

For better visualization of representative images shown, brightness and contrast were adjusted consistently across all samples and the images were converted to RGB.

#### ImageXpress acquisition and MetaXpress analysis

Labeled cells in 96-well flat bottom black/clear plates (Falcon 353219) were imaged with a 20×/0.45 NA objective on a Molecular Devices ImageXpress Micro XLS. For each well, twelve images (field of views; FOVs) were acquired and analyzed with Molecular devices MetaXpress software. The “Multiwavelength cell scoring” module was used to quantify the cell survival and also cell number for a particular marker. OPC outgrowth was measured using the “Neurite outgrowth” module, which quantifies the processes of cells. Data from the 12 images were averaged to a single data point per well, with seven well replicates per group. The number of surviving cells in each sample was then divided by the mean of the control samples to obtain the fold-change value. For better visualization of representative images shown, brightness and contrast were adjusted consistently across all samples.

#### Statistics

Microsoft Excel (Version 2022 Build 16.69.1) was used for collating data. All graphs were generated using GraphPad Prism 9.3.1. Shapiro-Wilk normality test was applied to verify normal distribution of data. One-way analysis of variance (ANOVA) with Tukey’s multiple comparison test was used to analyze statistically significant differences among multiple groups. Unpaired two-tailed Student’s *t*-tests was used to compare two groups. *P* < 0.05 was considered statistically significant. All values are shown as mean ± SEM.

## Results

### Upregulation of neurocan at perihematomal area and lesion core in murine ICH

The murine ICH lesions were assessed by Hematoxylin & Eosin (H&E) staining at day 3, 7 and 14 after collagenase-induced injury. We found the hematoma size to decrease over time (Supp. Fig. 1A), likely as a result of phagocytosis of erythrocytes and debris by microglia/macrophages and the resolution of edema as reviewed elsewhere.[4, 5] We chose day 7 after ICH for focus in this study, given a sizable lesion and also the accumulation of ECM molecules in the first week in other models of CNS injury.[20, 23] Also, at day 7 of murine ICH, there was prominent accumulation of Iba1^+^ microglia/macrophages and GFAP^+^ astrocytes, and NeuN^+^ neurons at the perihematomal area (Supp. Fig. 1B, 1C).

**Figure 1.**
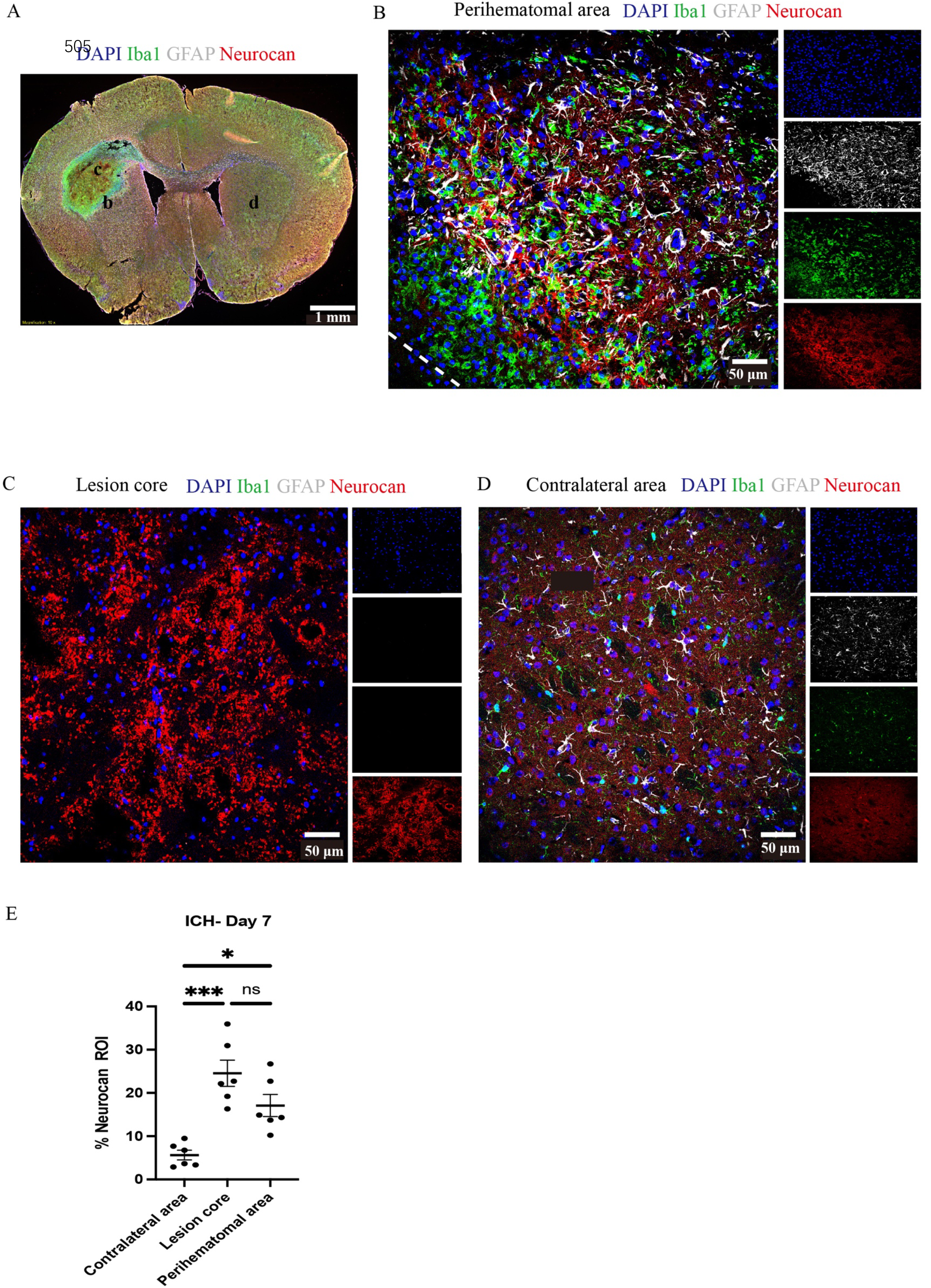
Neurocan is upregulated in perihematomal area and lesion core in murine ICH. **(A)** Representative slide scanner image of coronal brain sections from 7-day ICH mice shows DAPI for cell nuclei (blue), Iba1 for microglia/macrophages (green), GFAP for astrocytes (grey), and neurocan (red) in the perihematomal area (b), lesion core (c) or contralateral area (d). **(B-D)** Representative confocal images from 7-day ICH mice in perihematomal area **(B)**, lesion core **(C)** and contralateral area **(D)** stained for DAPI (blue), Iba1 (green), GFAP (grey), and neurocan (red). The individual colors are represented as individual boxes on the right while the merged image is to the left. The lower left corner inside the dotted lines in **B** is the lesion core. **(E)** Bar graphs comparing the levels of neurocan in perihematomal area, lesion core and contralateral area at day 7 of ICH, expressed as neurocan ^+^ in region of interest (ROI), where each ROI is a region defined by area occupied by Iba1^+^ cells (n=24 ROIs from 6 mice per group). Data are presented as the mean ± SEM, and analysed by one-way ANOVA-Tukey’s post hoc test; ns: not significant. Significance indicated as *P < 0.05, ***P < 0.001.

Next, we evaluated brain tissue sections at day 7 of ICH to address the relative elevation of CSPG members. The perihematomal area contained a high density of microglia/macrophages and reactive astrocytes, as reported by Iba1 and GFAP respectively. In the perihematomal area and lesion core (Fig. 1), but not in the contralateral uninjured hemisphere, neurocan immunoreactivity was prominent. Quantitative analysis shows that the elevation of neurocan was comparable between the perihematomal and lesion core of ICH (Fig. 1E).

No neurocan signal was observed when an isotype antibody control used in place of the primary rabbit neurocan antibody, or omission of neurocan antibody (secondary antibody control) was employed (Supp. Fig. 2).

**Figure 2.**
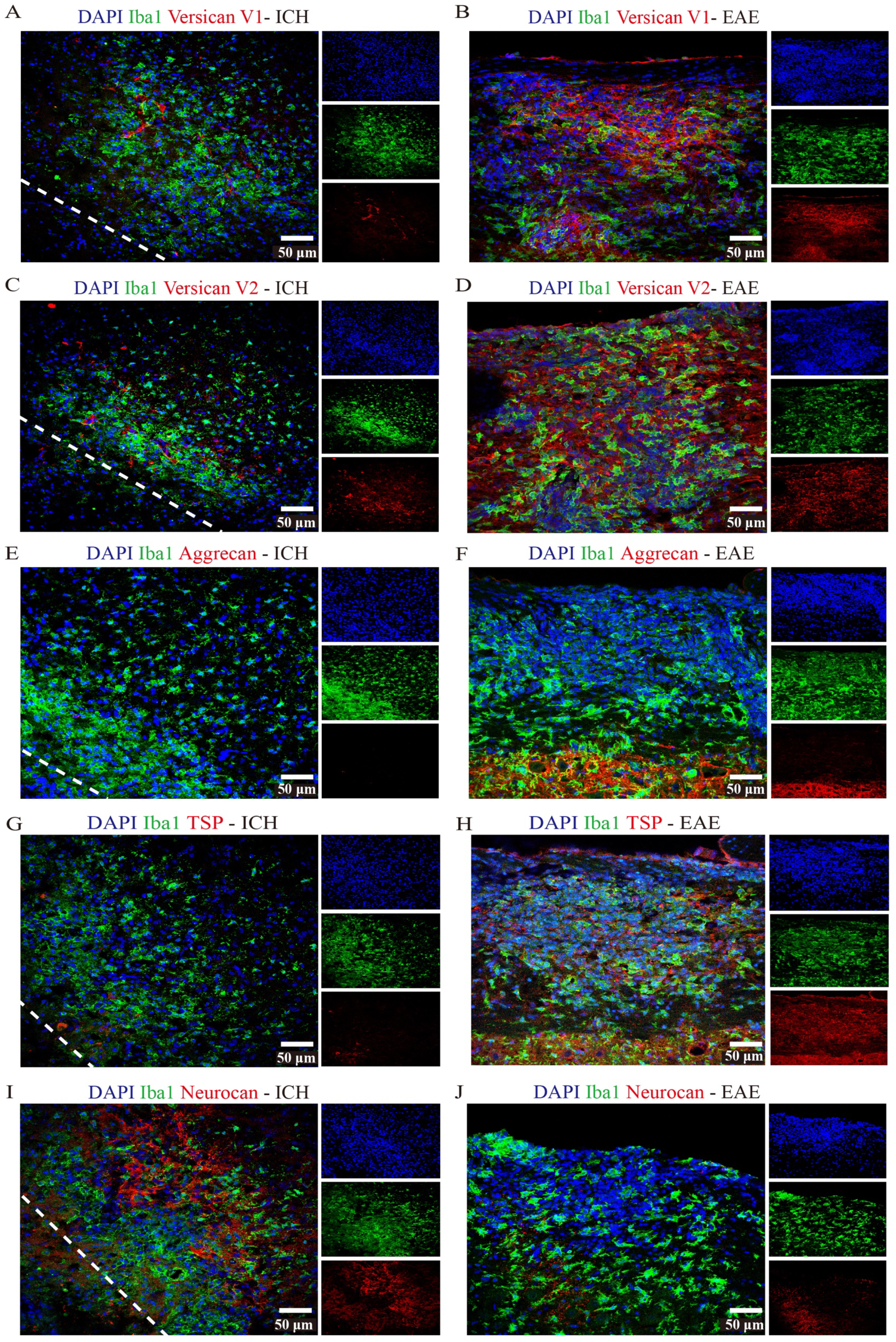
Non-neurocan CSPG members and TSP are minimal in ICH while robustly expressed in EAE lesions. Representative confocal images of sections from ICH mice at perihematomal area (left) and sections of spinal cord white matter from EAE mice (right). Stains are indicated by the respective colored labels. Images show versican-V1 **(A, B)**, versican-V2 (red) **(C, D)**, aggrecan (red) **(E, F)**, TSP (red) **(G, H)**, neurocan (red) **(I, J)**. Versican-V1, V2, TSP molecules are accumulated in EAE lesions but not in perihematomal area of ICH. The expression levels of aggrecan are low in both ICH and EAE lesions. However, neurocan molecules aggregate in the perihematomal area of ICH but not in EAE lesions. The lower left corner inside the dotted lines is the lesion core of ICH. Scale bar = 50 μm.

In contrast to neurocan upregulation, other lectican CSPG members were not detected following ICH. Immunoreactivity was not evident for versican-V1, versican-V2 or aggrecan in ICH, while these CSPGs were found in EAE tissue (Fig. 2). Further contrasting ICH and EAE, the high neurocan immunoreactivity in ICH tissue was negligible in EAE (Fig. 2J).

### Expression of other ECM components in murine ICH

TSP was not detected in ICH while present in EAE (Fig. 2G, H). Fibrinogen was present in the perihematomal and lesion core areas, albeit in higher amounts in the latter (Fig. 3A, D). This was also the case for fibronectin (Fig. 3B, E). HSPG was noted in the perihematomal area and lesion core (Fig. 3C, F), with a trend towards higher expression in the perihematomal area.

**Figure 3.**
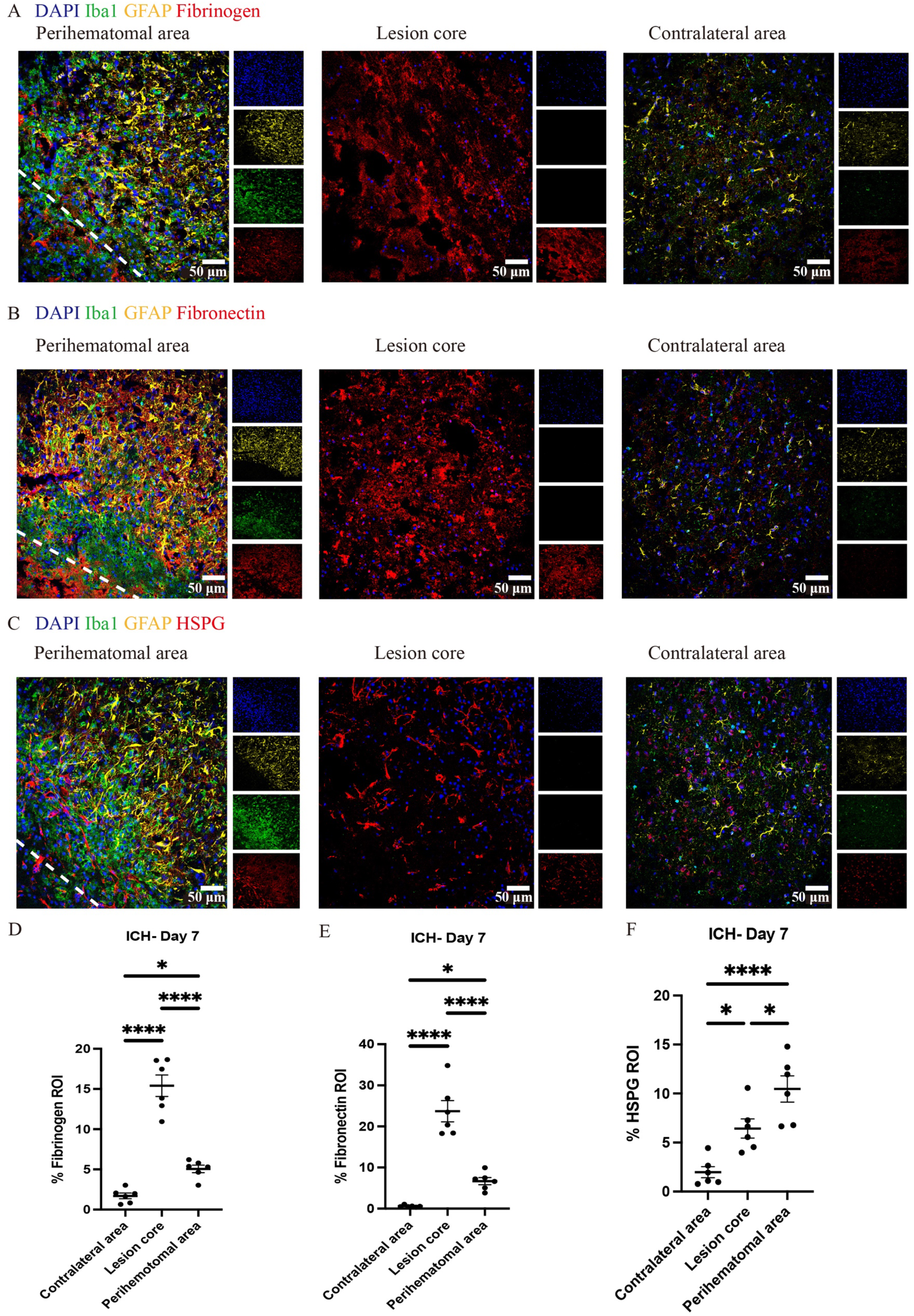
Fibrinogen, fibronectin and HSPG molecules are elevated at perihematomal area and lesion core in ICH mouse model. Representative confocal images of brain sections from ICH mice at perihematomal area, lesion core and contralateral area at day 7 stained for DAPI for cell nuclei (blue), Iba1 (green), GFAP (yellow), and fibrinogen (red) **(A)**, fibronectin (red) **(B)** or HSPG (red) **(C).** Scale bar = 50 μm. **(D-F)** Quantification shows the levels of ECM in perihematomal area and lesion core compared to contralateral area at day 7. N=6 replicates for each group; each dot represents mean of 4 locations of each area analyzed per mouse. Data are presented as the mean ± SEM. One-way ANOVA-Tukey’s post hoc test. Significance indicated as *P < 0.05, ****P < 0.0001.

### Cellular expression of ECM members that are elevated in murine ICH

Due to the high accumulation of neurocan, fibrinogen, fibronectin and HSPG in ICH, it was challenging to determine their precise localization within cell types. We thus utilized the 3D surface rendering of the Imaris program to better investigate the association of cells with these ECM molecules. The z-stack surface rendering of Iba1, GFAP and ECM molecules suggested that neurocan, fibrinogen, fibronectin and HSPG were intimately related intracellularly in Iba1^+^ and GFAP^+^ cells, as well as in proximity to these cells in the extracellular space (Fig. 4). There appears to be a higher internal accumulation of fibrinogen within GFAP^+^ compared to Iba1^+^ cells (Fig. 4B), and higher HSPG signal within Iba1^+^ compared to GFAP^+^ cells (Fig. 4F). Neurocan and fibronectin had a similar level of expression within GFAP^+^ and Iba1^+^ cells.

**Figure 4.**
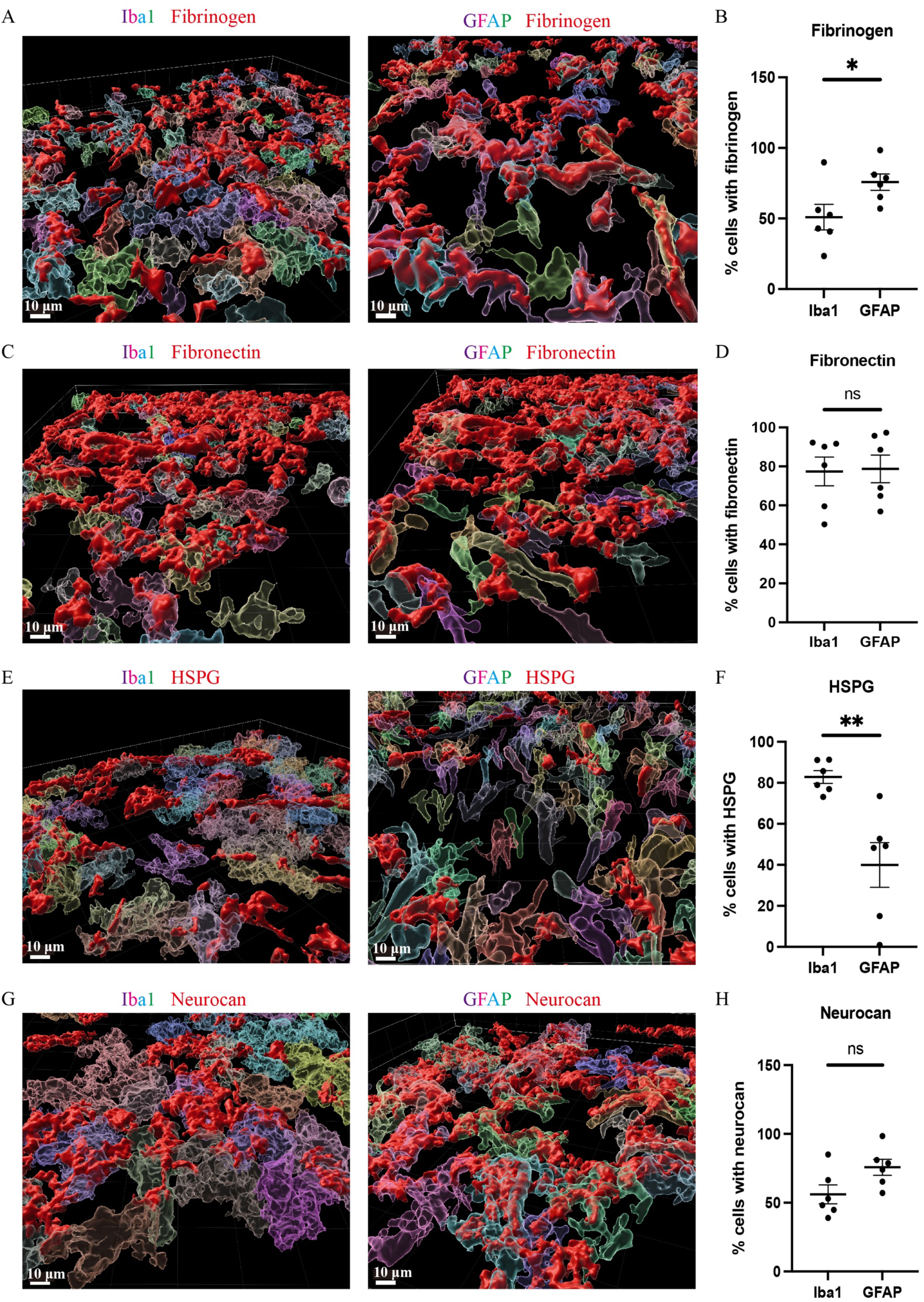
Fibrinogen, fibronectin, HSPG and neurocan within Iba1^+^ or GFAP^+^ cells. Representative 3D reconstruction images of perihematomal area at day 7 in ICH using Imaris rendition, internal accumulation of fibrinogen (red) **(A)**, fibronectin (red) **(C)**, HSPG (red) **(E)**, or neurocan (red) **(G)** within Iba1^+^ (left) or GFAP^+^ (right) cells. To determine the total count of Iba1^+^ or GFAP^+^ cells, as well as the number of Iba1^+^ or GFAP^+^ cells containing ECM molecules, each cell was individually labelled with a distinct color. Scale bar = 10 μm. We note that the fibrinogen, fibronectin, HSPG and neurocan staining is within Iba1^+^ or GFAP^+^ cells, and also in the extracellular space. **(B, D, F, H)** Quantification showing the percentage of Iba1^+^ or GFAP^+^ cells containing ECM molecules. Data are presented as the mean ± SEM of 6 mice. Each dot represents mean of 4 locations of each area analyzed per mouse. Unpaired two-tailed Student’s *t*-test; ns: not significant. Significance indicated as *P < 0.05, **P < 0.01.

### Neurocan inhibits OPCs in culture

Although ECM molecules can affect OPCs [7, 20], this is not known for neurocan. Given the prominence of neurocan expression in murine ICH, we evaluated its activity on OPCs. Figure 5 shows that when OPCs were seeded onto a substrate coated with neurocan, there were fewer O4+ OPCs that attached. Of the attached cells, these had fewer processes which, in vivo, are the precursors of myelin sheaths. Similarly, a mixed CSPG preparation used as a positive control also inhibited OPC adhesion and process outgrowth. Finally, fibronectin and fibrinogen were tested but these did not affect OPCs (Fig. 5).

**Figure 5.**
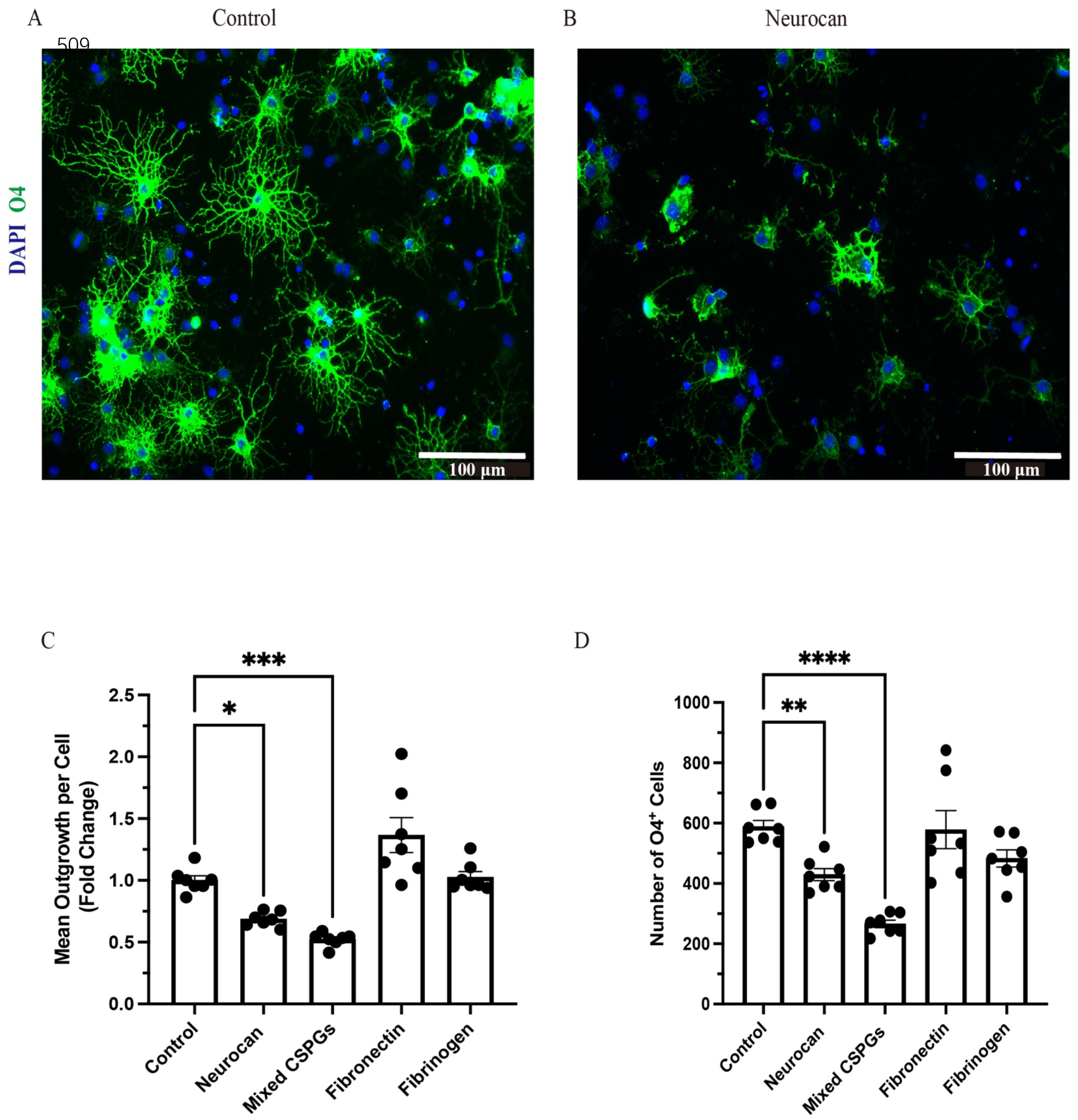
OPCs are inhibited by CSPGs and neurocan in culture. Representative images of mouse OPCs stained for the sulfatide O4 24h after plating onto PBS (control) **(A)** or neurocan **(B)** coated wells. Scale bar = 100 μm. Bar graphs comparing the fold change of mean process outgrowth of mouse OPCs **(C)**, or number of O4^+^ cells **(D)** cultured on control, neurocan, mixed CSPGs, fibronectin and fibrinogen after 24h (n=7 replicates). Data are presented as mean ± SEM. This was duplicated in two separate experiments. The single data point represents average of twelve images (field of views, FOVs) of each well of cells. One-way ANOVA with Dunnett post hoc; *P < 0.05, **P < 0.01, ***P < 0.001, ****P < 0.0001.

These results highlight that the neurocan elevation in murine could be a substantial impediment to OPCs that may attempt remyelination following ICH.

### High levels of neurocan in human ICH

We had the opportunity to corroborate the mouse data using a case of human ICH. The central hemorrhage core consisted purely of necrotic tissue and coagulated hemorrhage and could not be analysed. Two distinct penumbral and contralateral regions were analysed. Tissue sections were taken from hemorrhage periphery and perihematomal parenchyma (Fig. 6A), and the integrity of the tissue was informed by H&E staining (Fig. 6B). The latter highlighted diffuse parenchymal injury and hemorrhage in the affected frontal white matter. Importantly, similar to murine data, neurocan was highly deposited in perihematomal regions, closely associated with Iba1^+^ microglia/macrophages (Fig. 6C, 6E), as compared to contralateral region (Fig. 6D).

**Figure 6.**
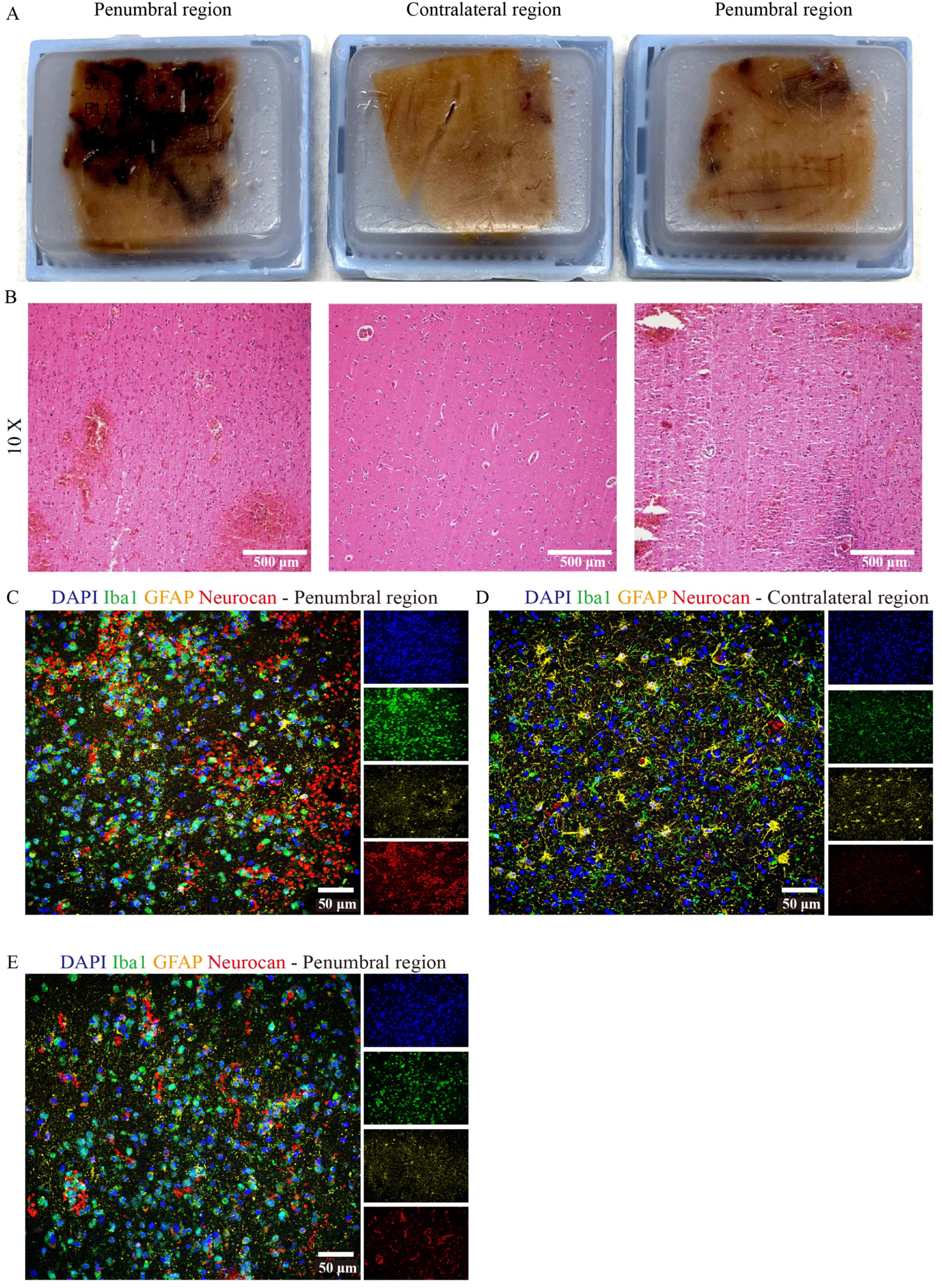
Neurocan is prominently increased in human ICH. **(A)** Human paraffin brain samples encompassing both hemorrhagic edge and perihematomal parenchyma regions, or the contralateral unaffected parenchyma, from a MCA infarct with hemorrhagic transformation. (**B)** Perihematomal and contralateral regions were stained with H&E to assess hemorrhage lesions. Scale bar = 200 μm. Representative immunofluorescent images of human brain tissue at different perihematomal **(C, E)** and contralateral **(D)** regions labeled with DAPI for cell nuclei (blue), Iba1 for microglia/macrophages (green), GFAP for astrocytes (yellow) and neurocan (red). Scale bar = 50 μm.

## Discussion

The poor prognosis of ICH is associated with mechanical disruption of brain by the enlarged hematoma and the progressive inflammatory responses.[4, 5, 27] Hematoma size gradually decreased over time (Supp. Fig. 1A), with persistent activated microglia/macrophages however, [28], and subsequently, the hematoma core became a cavity surrounded by GFAP^+^ astrocytes (Supp. Fig. 1C). While these events are well recognized in ICH, the deposition of ECM molecules is a dearth of knowledge in stroke.

Deposited ECM components play a significant role in inflammatory outcomes following CNS injury.[17] Findings from ischemic stroke studies have highlighted that specific ECM components including CSPGs, fibronectin, TSP, tenascin-C and laminins are elevated and exacerbate injury by enhancing neuroinflammation in the perihematomal area.[29-31] Similarly, there is an upregulation of aggrecan, brevican, neurocan and versican in rabbit pups after intraventricular hemorrhage.[21] CSPGs are known drivers of neuroinflammation, as well as inhibitors of remyelination and axonal regeneration in the CNS.[20, 32] Despite the pathologic significance of ECM, the expression and functions of ECM members in ICH have yet to be elucidated. Thus, we determined whether ECM molecules levels were altered in the collagenase-induced mouse model of ICH and in human ICH.

Our study is the first to investigate the changes of specific components of ECM in ICH. We found that neurocan was the predominant lectican CSPG in the perihematomal area and lesion core in ICH models, and expressed in perihematomal area of accumulated GFAP^+^ and Iba1^+^ cells (Fig. 1), consistent with previous studies showing astrocytes and macrophages are the major producers of CSPGs following non-ICH injury in the CNS.[7, 17, 33]. Neurocan was also highly accumulated in hemorrhagic and penumbral regions in human ICH (Fig. 6). In contrast, the levels of aggrecan, versican-V1 and versican-V2 molecules were not upregulated in ICH (Fig. 2). These results highlight the prominence of neurocan inside and around the hematoma in ICH. Strikingly, these results differ from those observed in EAE, with prominent versican-V1 but not neurocan elevation. Furthermore, TSP was not elevated in ICH while being highly expressed in EAE (Fig. 2). The reason and consequence of this differential expression deserves investigation in future studies.

We also found that fibrinogen, fibronectin and HSPG are upregulated in the perihematomal area and lesion core, and their expression in the perihematomal area are within GFAP^+^ and Iba1^+^ cells (Fig. 4). These findings are in accordance with other models in which fibronectin and HSPG molecules are expressed in CD45^+^ leucocytes, activated macrophages and reactive astrocytes following CNS injury.[7, 34] Transforming growth factor beta and epidermal growth factor signaling pathways contribute to the upregulation of ECM molecules in astrocytes.[35] The damage to the blood-brain barrier and deposition of plasma-derived ECM such as fibrinogen can also lead to their increase within the CNS parenchyma. These proteins have been shown to promote TGF-beta signaling in astrocytes, which can further stimulate the local production of ECM proteins.[36]

We observed that mixed CSPGs exert a more potent inhibitory effect on process outgrowth and differentiation of OPCs than neurocan (Fig. 5). One possible reason is that the CSPG mixture contains other lectican CSPG members including aggrecan, brevican and versican, which may bind to more receptors and act in synergy to inhibit OPCs.[37]

In summary, ICH is accompanied by elevation of several ECM molecules. One of these, neurocan, is a potent inhibitor of properties of OPCs in vitro and may hinder attempts at remyelination in vivo. The diversity of expressed ECM components in ICH suggests a multitude of functions in ICH that deserves future studies to define their distinct or collective roles in ICH. This study introduces the potential of targeting ECM as an untapped approach to improve the prognosis of ICH.

## Abbreviations

CNS: central nervous system
CSPGs: chondroitin sulphate proteoglycans
EAE: experimental autoimmune encephalomyelitis
ECM: extracellular matrix
GAG: glycosaminoglycan
GFAP: glial fibrillary acidic protein
HSPGs: heparan sulphate proteoglycans
Iba1: Ionized calcium-binding adapter molecule 1
ICH: intracerebral hemorrhage
MCA: middle cerebral artery
MMPs: matrix metalloproteinases
OPC: oligodendrocyte precursor cells
PBS: Phosphate-buffered saline
PFA: Paraformaldehyde
TSP: thrombospondin.

## Acknowledgements

We thank the Hotchkiss Brain Institute Advanced Microscopy Platform facility for providing microscopy and image analysis platforms. We acknowledge operating grant support from the Canadian Institutes of Health Research (Foundation grant 1049959) (VWY); and from the National Key Research and Development Program of China (grant no: 2018YFC1312200), and the National Natural Science Foundation of China (grants no: 82071331, 81870942, and 81520108011) (MX). HL is supported by a PhD studentship from the China Scholarship Council.

## Competing interests

The authors declare no potential conflicts of interest with respect to the research, authorship and/or publication of this article.

## Notes

### Competing Interest Statement

The authors have declared no competing interest.

