## Supplemental Figure for "Prominent elevation of extracellular matrix molecules in intracerebral hemorrhage"

**Supplementary Figure S1. ICH injury in mice induced by collagenase.**

**(A)** Frozen brain sections from ICH mice at different time points were stained with H&E to identify lesions. Black rings indicate the hematoma sites. Scale bar = 1mm. **(B)** Representative images of coronal brain sections from ICH mice labeled with DAPI for cell nuclei (blue), Iba1 (green), GFAP (grey) and NeuN (red) showing the site of the perihematomal area (c). Scale bar = 1mm. **(C)** Representative confocal images of brain sections from ICH mice at perihematomal area at day 7 stained for DAPI (blue), Iba1 (green), GFAP (grey) and NeuN (red). The lower left corner within the dotted lines is the lesion core. Scale bar = 50 μm.

**Supplementary Figure S2. Isotype control and secondary antibody control of neurocan staining.**

**(A)** Representative confocal images of brain sections from ICH mice at perihematomal area, lesion core and contralateral area at day 7 stained with DAPI for cell nuclei (blue), Iba1 (green) and neurocan (red). **(B)** Representative confocal images of brain sections from ICH mice at perihematomal area, lesion core and contralateral area at day 7 stained with DAPI for cell nuclei (blue), Iba1 (green) and isotype antibody (red). **(C)** Representative confocal images of brain sections from ICH mice at perihematomal area, lesion core and contralateral area at day 7 stained with only secondary antibodies and DAPI (blue). The lower left corner within the dotted lines is the lesion core. Scale bar = 50 μm.

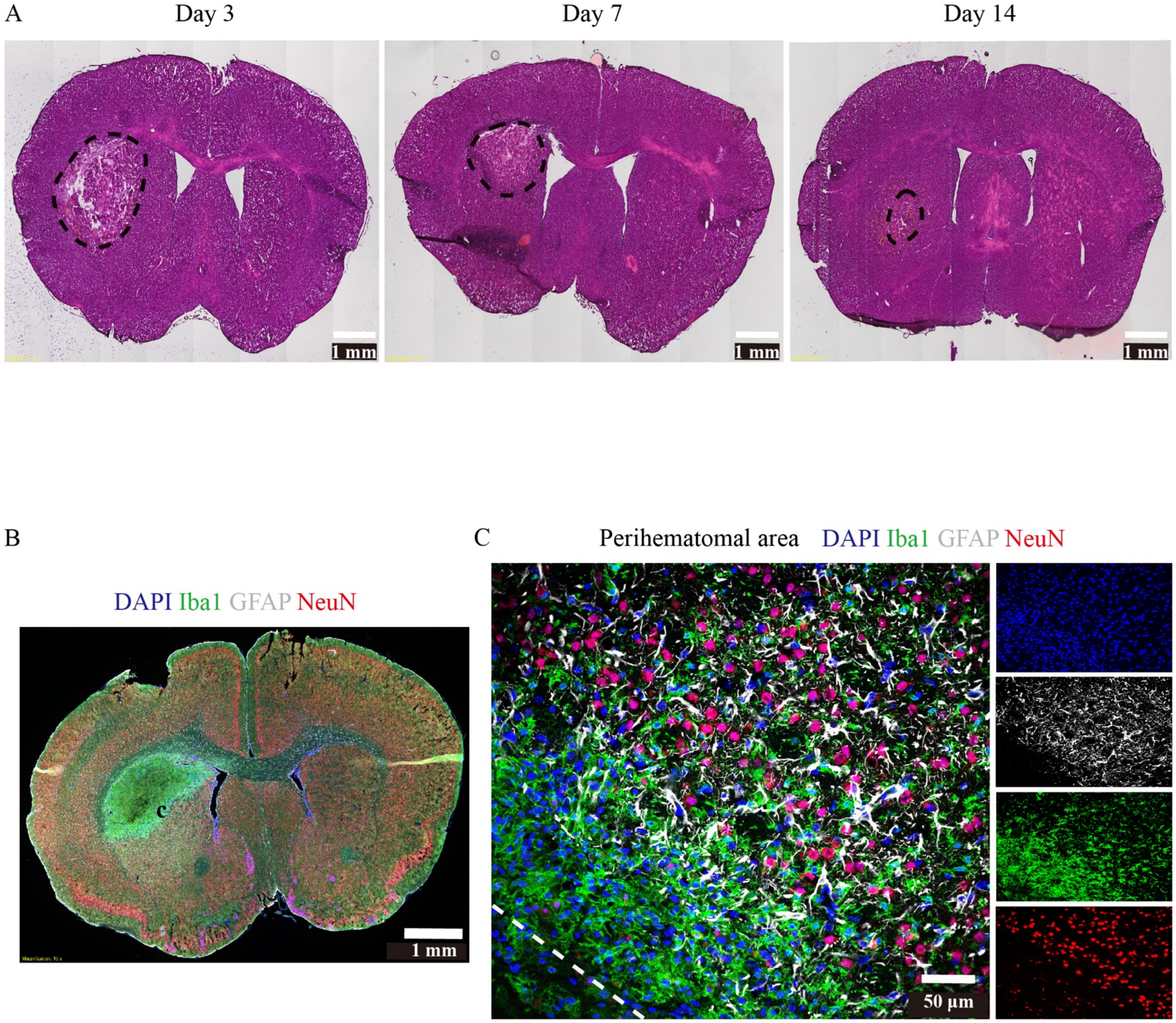
A
Day 3
Day 7
Day 14
1 mm
1 mm
1 mm
B
DAPI Iba1 GFAP NeuN
c
1 mm
C
Perihematomal area
DAPI Iba1 GFAP NeuN
50 μm

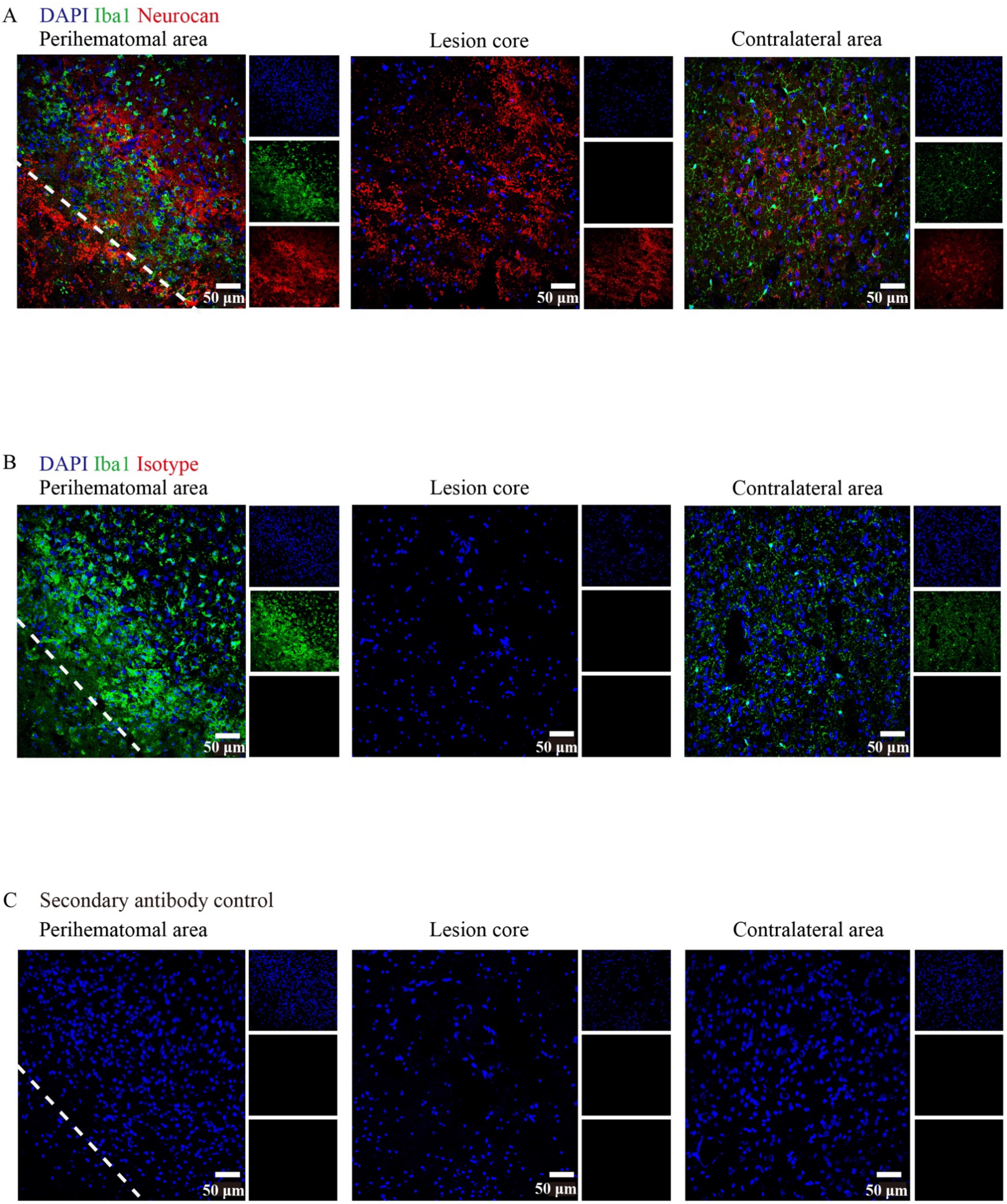
A
DAPI Iba1 Neurocan
Perihematomal area
Lesion core
Contralateral area
50 µm
B
DAPI Iba1 Isotype
Perihematomal area
Lesion core
Contralateral area
50 µm
C
Secondary antibody control
Perihematomal area
Lesion core
Contralateral area
50 µm
